## Supplementary Information for "A molecularly defined basalo-prefrontal-thalamic circuit regulates sensory and affective dimensions of pain in male mice"

### Supplementary Figures and legends

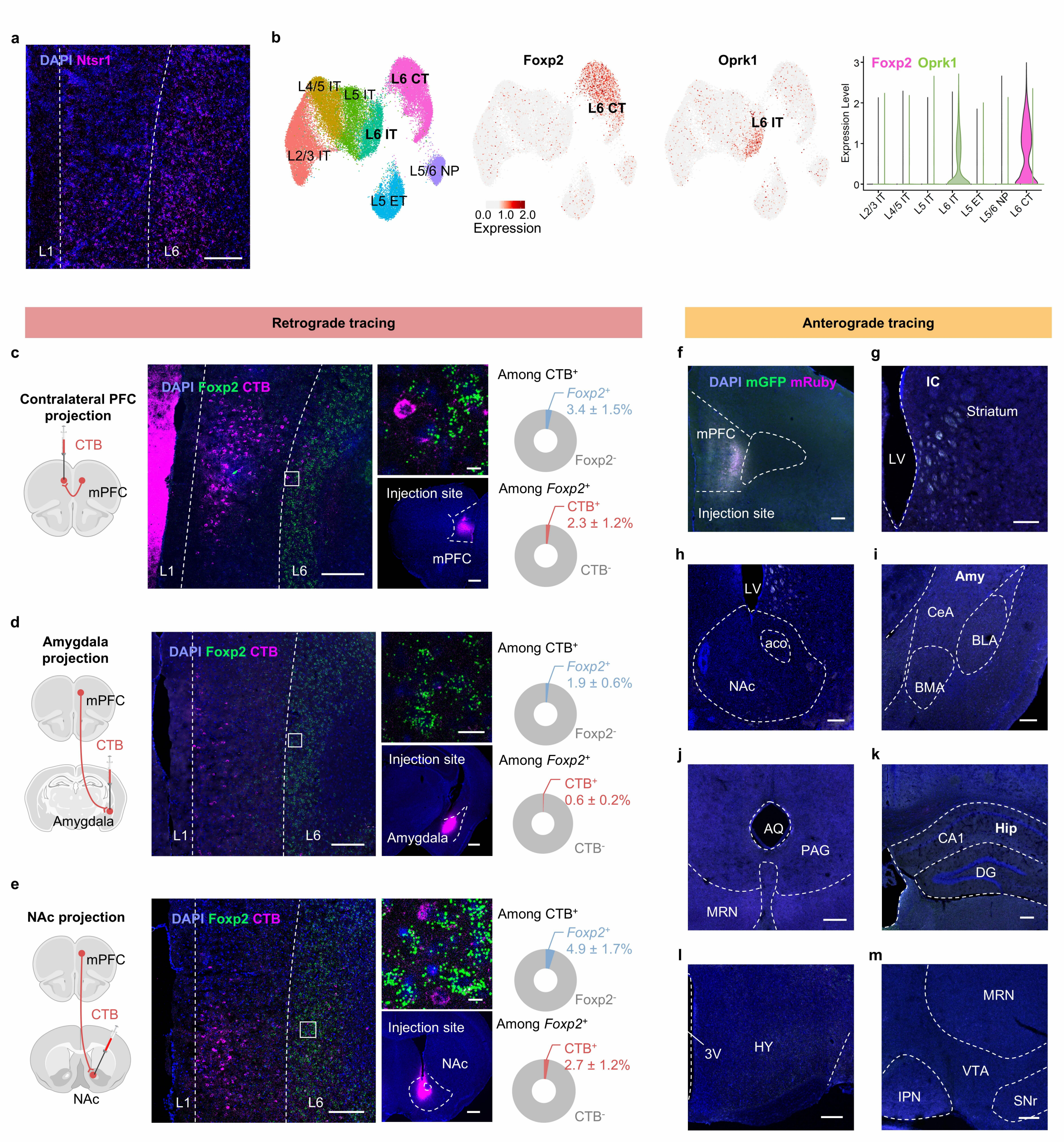


**Supplementary Fig. 1. *Foxp2* specifically marks the L6 corticothalamic neurons in the mPFC.** **a**, smFISH for *Ntsr1* (magenta) in the mPFC. Scale bar: 200 μm. n = 3 mice. **b**, UMAP showing the results of an integrated analysis of single-cell RNA sequencing and spatial transcriptomics with neuron types and layer information (left). Predicted expression patterns of *Foxp2* (middle left) and

*Oprk1* (middle right). L, layer; CT, corticothalamic neurons; ET, extratelencephalic neurons; IT, intratelencephalic neurons; NP, near-projecting neurons. Predicted expression levels of *Foxp2* and *Oprk1* in the mPFC neuronal subtypes with layer information (right). **c**, Diagram of CTB injection into the contralateral mPFC (cPFC) (left). Representative image of CTB (magenta) and smFISH for *Foxp2* (green) in the mPFC (middle), enlarged image of the indicated region (middle top) and the injection site of CTB (middle bottom). Right, quantification of the $CTB^+$ neurons co-expressing *Foxp2* (top right) or the *Foxp2*$^+$ neurons co-labeled by CTB (bottom right). Scale bars: middle left, 200 μm; middle top, 10 μm; middle bottom, 500 μm; n = 3 mice. **d**, **e**, Same as **c** but for CTB injected into the amygdala (**d**) or NAc (**e**). n = 3 mice. **f**, Representative image of virus expression in the mPFC of *Foxp2*-Cre mice as indicated in the **Fig. 1h**. Scale bar: 200 μm. **g-m**, The mGFP and synaptophysin-fused mRuby signals labeling the axon puncta of the mPFC *Foxp2*$^+$ neurons in the known targets of the mPFC, including striatum (**g**), NAc (**h**), amygdala (**i**), PAG (**j**), HPC (**k**), HY (**l**) and VTA (**m**). Scale bars: 200 μm. n = 3 mice. Data are represented as mean ± SEM. 3V, third ventricle; AQ, cerebral aqueduct; BMA, basomedial amygdala; CeA, central amygdala; DG, dentate gyrus; Hip, hippocampus; HY, hypothalamus; IPN, interpeduncular nucleus; LV, lateral ventricle; MRN, midbrain reticular nucleus; NAc, nucleus accumbens; SNr, substantia nigra, reticular part; VTA, ventral tegmental area. Source data are provided as a Source Data file. Created in BioRender. Liu, Y. (2026) https://BioRender.com/jpcdwkx.

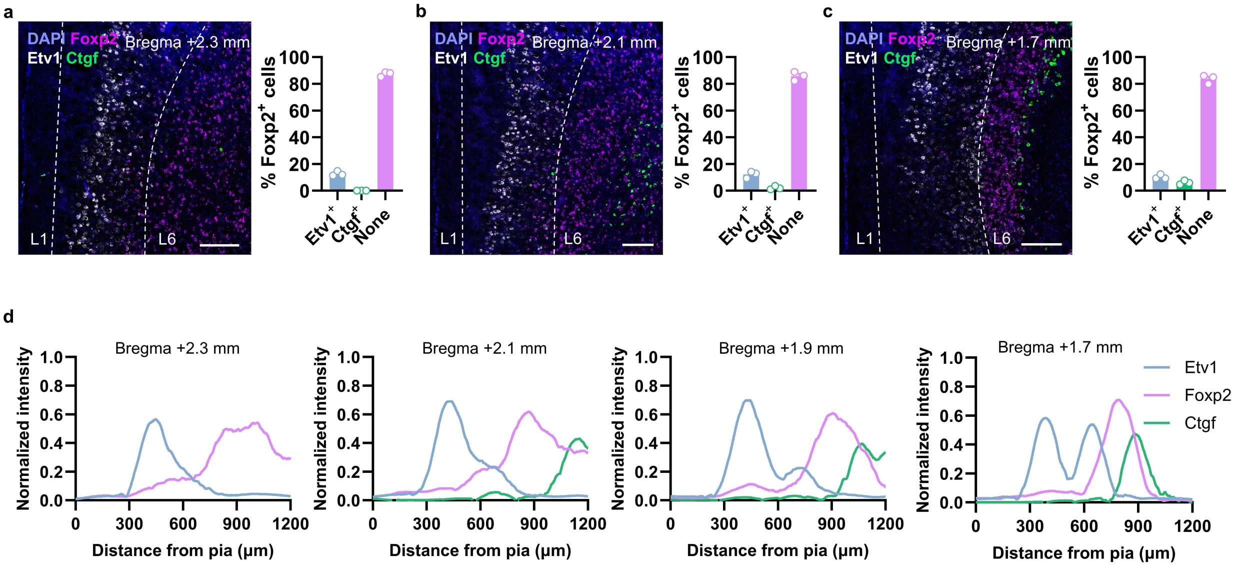


**Supplementary Fig. 2. Expression pattern of *Foxp2* in the mPFC along the anterior-posterior axis.**

**a**, smFISH for *Foxp2* (magenta), *Etv1* (white) and *Ctgf* (green) in the mPFC at 2.3 mm anterior to the bregma (left) and the percentages of the *Foxp2*$^{+}$ neurons co-expressing *Etv1* or *Ctgf* (right). Scale bar, 200 μm. n = 3 mice. **b**, **c**, Same as **a** but for the mPFC region 2.1 mm (**b**) or 1.7 mm (**c**) anterior to the bregma. n = 3 mice. **d**, Quantification of the normalized fluorescence signals of *Etv1* (blue), *Foxp2* (magenta) and *Ctgf* (green) in the mPFC from the pia to the white matter at the indicated sections of mPFC along the anterior-posterior axis. n = 3 mice. Data are represented as mean ± SEM. Each dot indicated one mouse. Source data are provided as a Source Data file.

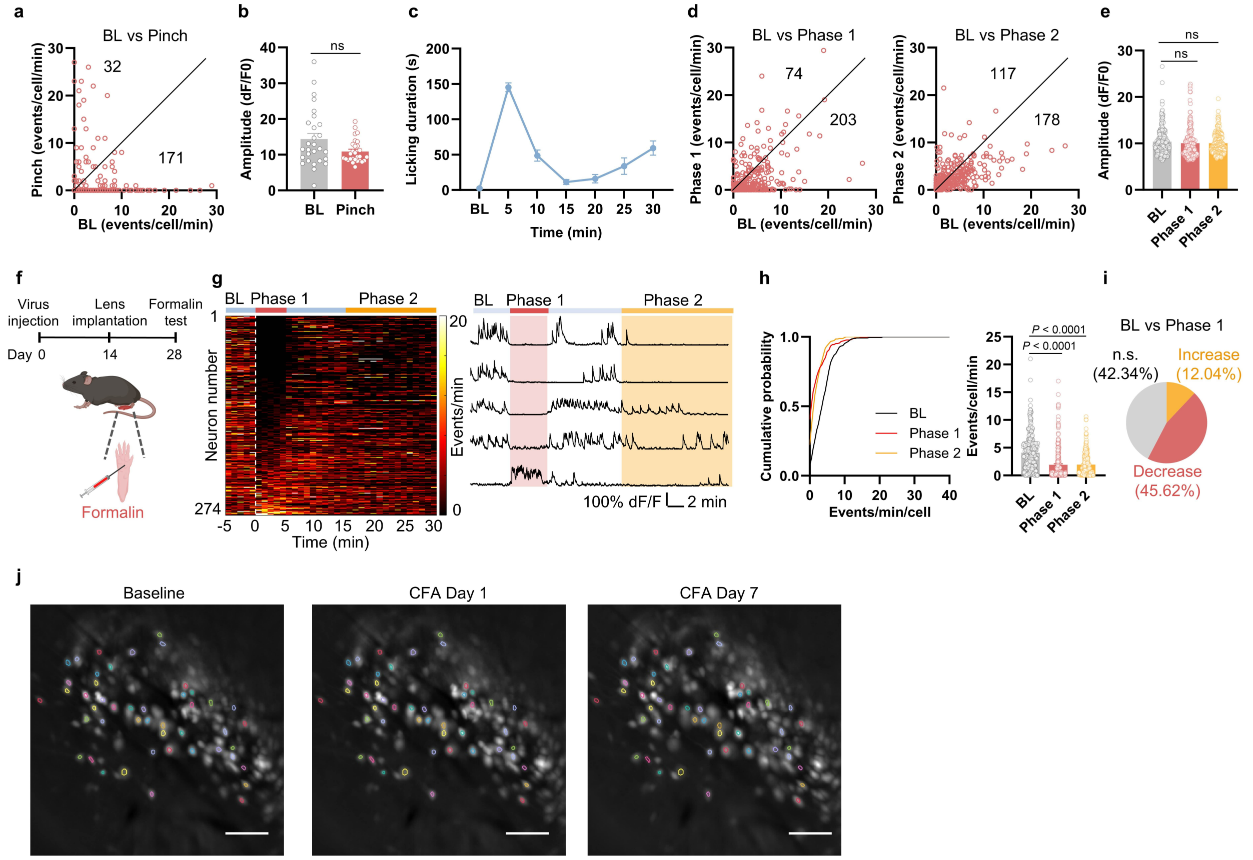


**Supplementary Fig. 3.** ***In vivo*** **miniscopic calcium imaging of the mPFC** ***Foxp2***$^{+}$ **neurons.**

**a**, Calcium event frequency of the identified neurons during baseline (BL) and pinch stimulation. The numbers of the neurons with increased or decreased frequency are indicated. n = 248 neurons from 5 mice. **b**, Amplitude of the calcium events during BL and pinch stimulation. Only the neurons that exhibited calcium events during both sessions were included (n = 34 neurons from 5 mice). Two-tailed Wilcoxon matched-pairs signed rank test. **c**, Licking duration of the mice subjected to formalin injection. n = 5 mice. **d**, Same as **a** but for formalin treatment comparing BL with phase 1 (left) or BL with phase 2 (right). n = 302 neurons from 5 mice. **e**, Same as **b** but for formalin treatment. Only the neurons that exhibited calcium events during all sessions were included. n = 196 neurons from 5 mice. RM one-way ANOVA with Sidak's multiple comparison, $F_{1.637,319.3} = 2.252$. **f**, Diagram of the miniscopic calcium imaging for formalin treatment without prior pinch or CFA treatment. **g**, Left, heatmap of the calcium event frequency averaged to 1 min of bin before and after formalin treatment (n = 274 neurons from 3 mice). The white dashed line indicates the start of formalin injection. Right, representative traces of calcium signals (ΔF/F) of individual neurons. The red and the yellow backgrounds indicate the periods of phase 1 and phase 2, respectively. **h**, Cumulative distribution (left) and the average (right) of calcium event frequency in baseline (BL), phase 1 and phase 2. n = 274 neurons from 3 mice. RM one-way ANOVA with Tukey's multiple comparison, $F_{1.809,493.7} = 67.21$. **i**, Percentages of the neurons with significantly decreased (red), increased (yellow), or not significant (n.s.) change (grey) in calcium event frequency during phase 1 compared to baseline. **j**, Representative field of view images in baseline, day 1 and day 7 after CFA injection. Some neuronal contours in the same locations are identified in different days and longitudinally registered as the same individual neurons, which are labeled in the images. Scale bars: 100 μm. n = 5 mice. Data are represented as mean ± SEM. ns, no significant difference. Source data are provided as a Source Data file. Created in BioRender. Liu, Y. (2026) https://BioRender.com/jpcdwkx.

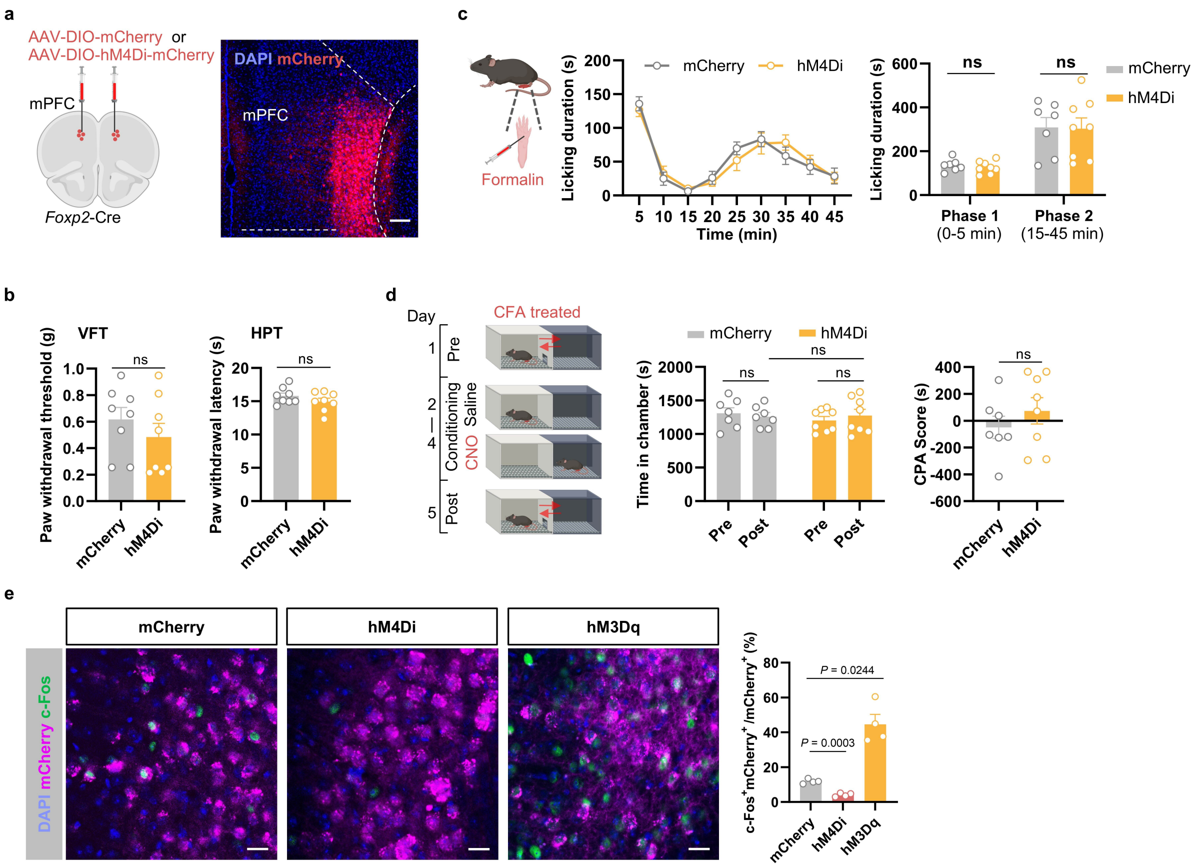


**Supplementary Fig. 4. Effects of inactivating the mPFC *Foxp2*$^+$ neurons on pain-related behaviors.**

**a**, Diagram of virus injection (left) and a representative image of hM4Di-mCherry expression in the mPFC of *Foxp2*-Cre mice (right). Scale bar: 100 µm. **b**, Paw withdrawal threshold and latency of the mice assessed by VFT (left) and HPT (right). n = 8 mice for each group. Two-tailed unpaired *t*-test, VFT, $t_{14}$ = 0.9739; HPT, $t_{14}$ = 1.241. **c**, Diagram of formalin test (left). Licking duration of the mice in response to formalin injection after CNO treatment (middle) and summarized licking duration in the phase 1 and phase 2 (right). n = 7 mice for mCherry group and 8 mice for hM4Di group. RM two-way ANOVA with Sidak's multiple comparison, $F_{1,13}$ = 0.04740. **d**, Diagram of the conditioned place avoidance (CPA) test (left). The time that mice spent in the chamber paired with CNO treatment (middle) and CPA scores (right) of the mice in CPA test (animal number is same as in **c**). Left, RM two-way ANOVA with Sidak's multiple comparison, $F_{1,13}$ = 0.2883; Right,

Two-tailed unpaired *t*-test, $t_{13} = 0.9380$. **e**, Representative images of the c-Fos expression in the mPFC after CNO treatment in the mCherry-expressing, hM4Di-mCherry-expressing or hM3Dq-mCherry-expressing mice. Right: percentage of the c-Fos$^{+}$ cells in the mPFC virus-expressing neurons. Scale bar: 20 μm. n = 4 mice for each group. Brown-Forsythe's one-way ANOVA test with Dunnett's T3 multiple comparison, $F_{2,3.138} = 42.64$. Data are represented as mean ± SEM. ns, no significant difference. Source data are provided as a Source Data file. Created in BioRender. Liu, Y. (2026) https://BioRender.com/jpcdwkx.

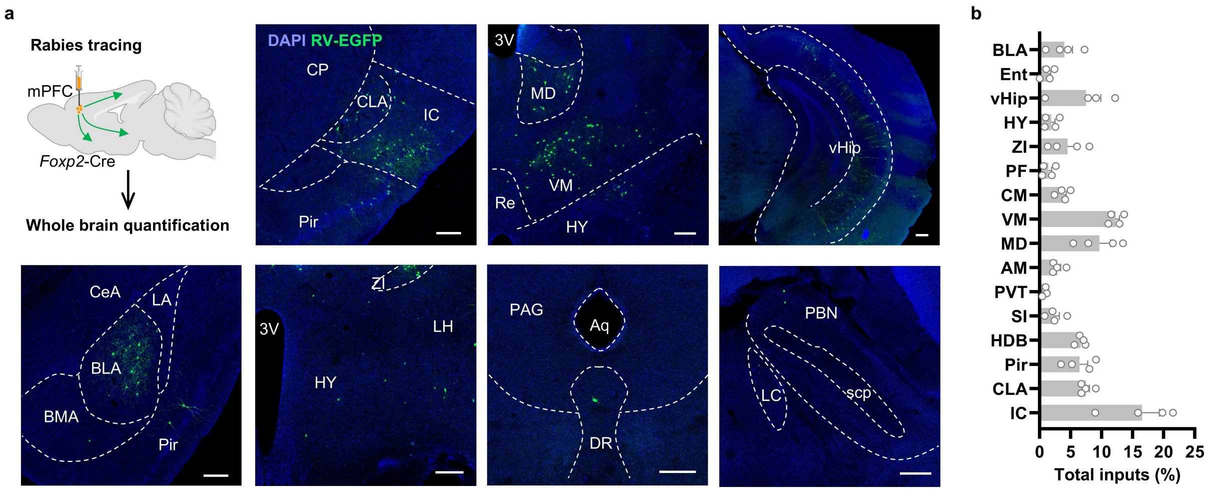


**Supplementary Fig. 5. Inputs of the mPFC *Foxp2*$^+$ neurons.**

**a**, Monosynaptic retrograde rabies tracing of the mPFC *Foxp2*$^+$ neurons (top left) and whole brain mapping. Representative images of brain regions with RV-EGFP labeled cells, including IC (top, 2$^{nd}$), TH (top, 3$^{rd}$), Hip (top, 4$^{th}$), BLA (bottom left), HY (bottom, 2$^{nd}$), PAG (bottom, 3$^{rd}$) and PBN (bottom, 4$^{th}$). Scale bars: 200 µm. n = 4 mice. **b**, Quantification of the retrogradely labeled cells in different brain regions. n = 4 mice. Data are represented as mean ± SEM. 3V, third ventricle. Aq, cerebral aqueduct; BLA, basolateral amygdala; BMA, basomedial amygdalar nucleus; CeA, central amygdalar nucleus; CP, caudoputamen; CLA, claustrum; DR, dorsal raphe nucleus; HDB, horizonal diagonal band of Broca; HY, hypothalamus; IC, insula cortex; LA, lateral amygdalar nucleus; LC, locus ceruleus; LH, lateral hypothalamic area; LPO, lateral preoptic area; MA, magnocellular nucleus; MD, mediodorsal thalamus; OT, olfactory tube; PAG, periaqueductal gray; PBN, parabrachial nucleus; Pir, piriform area; Re, nucleus of reuniens; scp, superioi cerebellar pedunles; SI, substantial insominata; TH, thalamus; vHip, ventral hippocampus; VM, ventromedial thalamus; ZI, zona incerta. Source data are provided as a Source Data file. Created in BioRender. Liu, Y. (2026) https://BioRender.com/jpcdwkx.

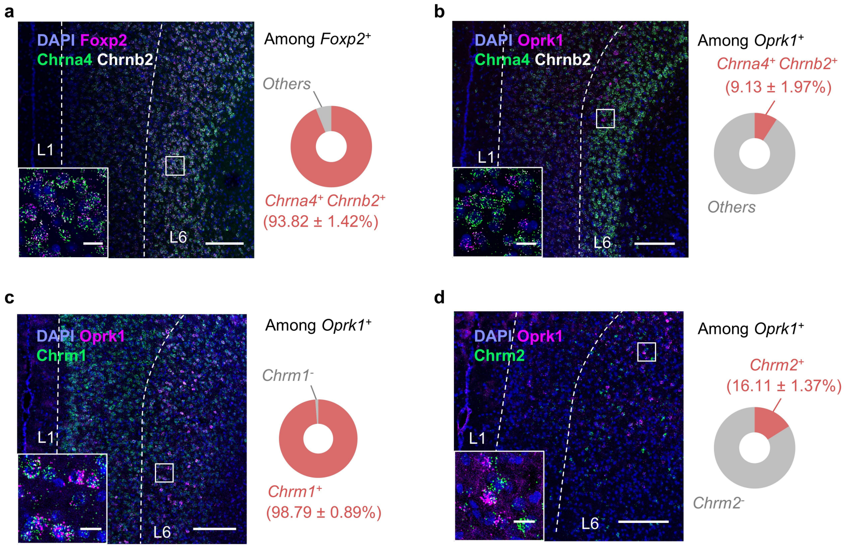


**Supplementary Fig. 6. Cell type-specific expression patterns of acetylcholine receptor in the mPFC.**

**a**, Left, representative images of smFISH experiments in the mPFC for *Foxp2* (magenta), *Chrna4* (green) and *Chrnb2* (white). An enlarged image of the indicated region is inserted in the bottom left box. Right: Quantification of $Chrna4^+Chrnb2^+Foxp2^+$ cells among all the $Foxp2^+$ cells. Scale bars: 200 μm; inserted box, 20 μm. n = 3 mice. **b**, Left: same as **a** but for *Oprk1* (magenta), *Chrna4* (green) and *Chrnb2* (white). Right: Quantification of $Chrna4^+Chrnb2^+Oprk1^+$ cells among all the $Oprk1^+$ cells. n = 3 mice. **c**, Left: smFISH of *Oprk1* (magenta) and *Chrm1* (green) in the mPFC. Right: quantification of the $Chrm1^+Oprk1^+$ cells among all the $Oprk1^+$ cells. Scale bars: 200 μm; inserted box, 20 μm. n = 3 mice. **d**, Same as **c** but for *Oprk1* (magenta) and *Chrm2* (green) in the mPFC. n = 3 mice. Source data are provided as a Source Data file.

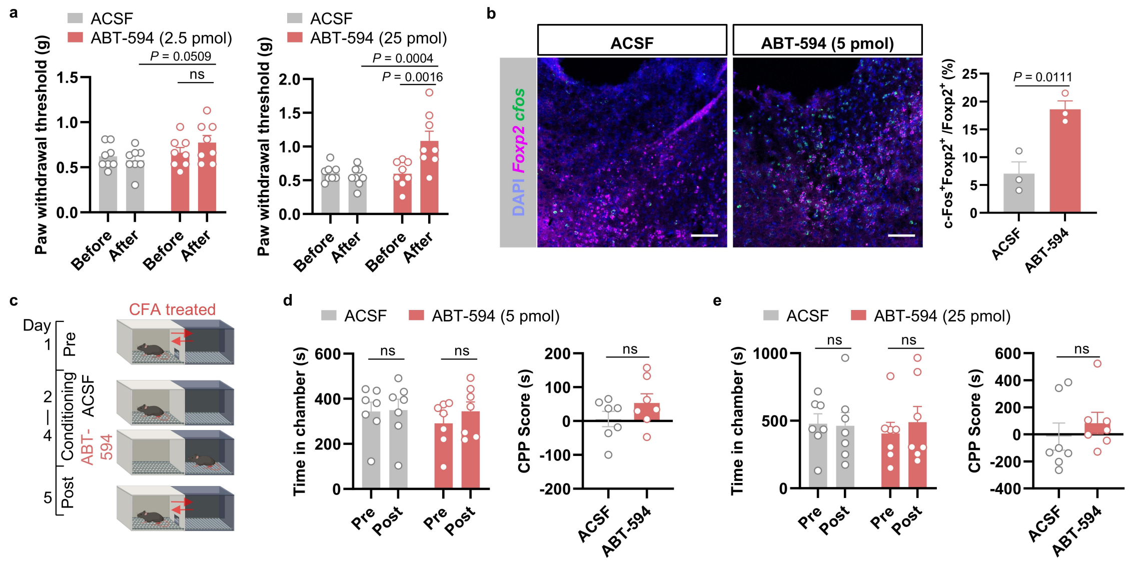


**Supplementary Fig. 7. Effects of activating the α4β2 nAChR in the mPFC.**

**a**, Dosage test of ABT-594. Paw withdrawal threshold of the mice assessed in VFT before and after the intracranial injection of ABT-594 (2.5 pmol/site) (left) or ABT-594 (25 pmol/site) (right) into the mPFC compared to ACSF treatment. n = 8 mice for each group. RM two-way ANOVA with Sidak's multiple comparison, $F_{1,14}$ = 6.077 (left), $F_{1,14}$ = 7.914 (right). **b**, Left, representative images of smFISH for *Foxp2* (magenta) and *cfos* (green) in mPFC after drug treatment. Right, percentage of the *cfos*$^+$ cells among the mPFC *Foxp2*$^+$ cells. Scale bars, 100 μm. n = 3 mice. Two-tailed unpaired *t*-test, $t_4$ = 4.461. **c**, Diagram of CPP test after CFA treatment paired with ACSF or ABT-594 treatment. **d**, Time spent in the chamber before and after training (left) and CPP scores (right) for ABT-594 treatment at 5 pmol per site. n = 7 mice for each group. **e**, Same as **d** but for ABT-594 treatment at 25 pmol per site. n = 7 mice for each group. RM two-way ANOVA with Sidak's multiple comparison for **d** (left, $F_{1,12}$ = 0.2535) and **e** (left, $F_{1,12}$ = 0.03665); two-tailed unpaired *t*-test for **d** (right, $t_{12}$ = 1.358) and **e** (right, $t_{12}$ = 0.7715). Data are represented as mean ± SEM. ns, no significant difference. Source data are provided as a Source Data file. Created in BioRender. Liu, Y. (2026) https://BioRender.com/jpcdwkx.
